# Radula diversification promotes ecomorph divergence in an adaptive radiation of freshwater snails

**DOI:** 10.1101/2020.01.17.910034

**Authors:** Leon Hilgers, Stefanie Hartmann, Jobst Pfaender, Nora Lentge-Maaß, Thomas von Rintelen, Michael Hofreiter

**Affiliations:** Institute of Biochemistry and Biology, University of Potsdam, Potsdam, Germany; Museum für Naturkunde Berlin, Leibniz Institute for Evolution and Biodiversity Science, Berlin, Germany; Zoological Research Museum Alexander Koenig, Leibniz Institute for Animal Biodiversity, Bonn, Germany; Naturkundemuseum Potsdam, Potsdam, Germany; Centrum für Naturkunde, Hamburg, Germany

## Abstract

Adaptive diversification of complex traits plays a pivotal role for the evolution of organismal diversity. However, the underlying molecular mechanisms remain largely elusive. In the freshwater snail genus *Tylomelania,* adaptive radiations were likely promoted by trophic specialization via diversification of their key foraging organ, the radula. To investigate the molecular basis of radula diversification and its contribution to lineage divergence, we use pooled tissue-specific transcriptomes of two sympatric *Tylomelania sarasinorum* ecomorphs. We show that divergence in both gene expression and coding sequences is stronger between radula transcriptomes compared to mantle and foot transcriptomes. These findings support the hypothesis that diversifying selection on the radula is driving speciation in *Tylomelania* radiations. We also identify several candidate genes for radula divergence. Putative homologs of some candidates (*hh*, *arx*, *gbb*) also contributed to trophic specialization in cichlids and Darwin’s finches, indicating that some molecular pathways may be especially prone to adaptive diversification.

## Main

Adaptive radiations provide extreme examples of rapid phenotypic and ecological diversification and therefore feature prominently among model systems for adaptation and speciation^1–6^. In many adaptive radiations, lineage divergence is promoted by diversification of a few traits, like foraging organs, which acted as key adaptive traits in several radiations^3,7–13^. Understanding the genetic bases of key adaptive traits is essential because they shape evolutionary trajectories of diversifying lineages^14,15^. Although previous findings are likely biased towards few genes of large effect^2,16^, they also indicate that polygenic selection^17–19^, adaptive introgression^20–24^, and regulatory evolution^18,21,25,26^ promote diversification in adaptive radiations^17–19^. However, much remains to be discovered about the genetic basis of adaptive traits, the molecular evolution underlying their diversification, and their contribution to speciation^2,27^. Particularly, our understanding of gene expression divergence and its contribution to speciation is still in its infancy^28,29^. Here we investigate the genetic basis of diversification of the molluscan key foraging organ (the radula) and its role for lineage divergence in a radiation of freshwater snails, using two sympatric ecomorphs of *Tylomelania sarasinorum*^30^.

The genus *Tylomelania* is endemic to the central Indonesian island Sulawesi and underwent several radiations following colonizations of different lake systems^31,32^. Lacustrine species flocks occur across heterogeneous substrates and exhibit remarkable radula diversity (Figure 1)^33,34^. In contrast, riverine clades occupy relatively homogenous substrates, have uniformly shaped radular teeth and include comparatively few species^33,34^. Additionally, similar radula morphologies likely evolved several times on similar substrates in different lakes^31,32,34^. Hence, it was hypothesized that divergent adaptation of the radula allowed efficient foraging on alternative substrates and promoted speciation in radiations of *Tylomelania*^31,32^.

**Fig. 1:**
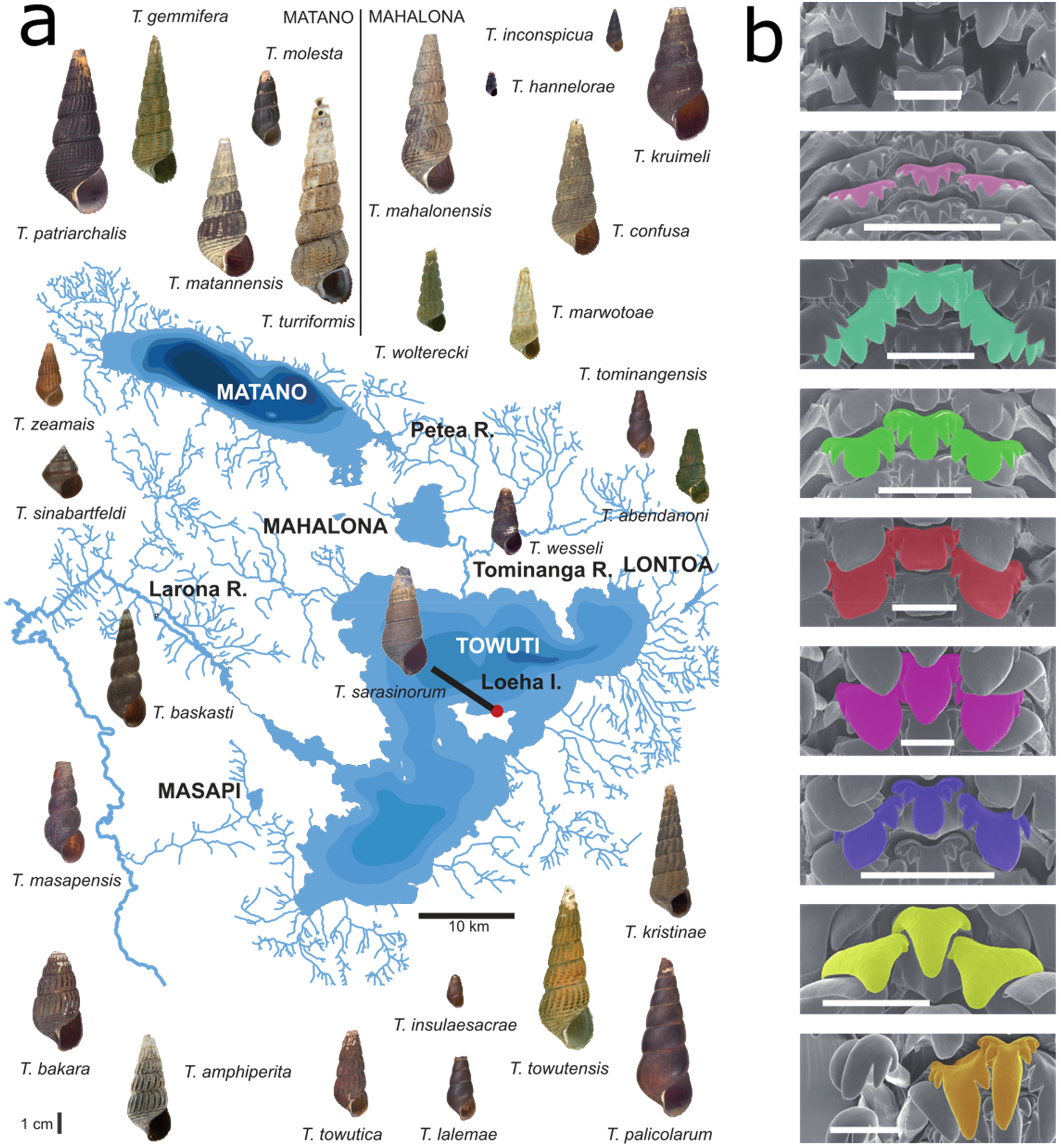
Diversity of the viviparous freshwater snail genus *Tylomelania* in the Malili Lakes System. Species diversity in the Malili Lakes and surrounding rivers (a) is shown together with an overview of radula morphologies (b) (Scale bars = 0.1 mm). The sampling site of *T. sarasinorum* specimens at Loeha Island is indicated by a red dot. It was hypothesized that ecomorphs of *T. sarasinorum* modified their radula in adaption to different microhabitats, i.e. submerged logs and rocks. Modified with permission from ^31^ and ^32^.

In addition to interspecific variation, some species exhibit radula polymorphisms^34^. One such species is *Tylomelania sarasinorum*, which reportedly has a substrate-correlated radula polymorphism. Ecomorphs occur on rocks and logs in shallow waters of Lake Towuti (Figure 1, 2a)^34^, but cannot be distinguished based on mitochondrial markers^31,35^. Given the radula’s hypothesized role as key adaptive trait in this radiation, *T. sarasinorum* ecomorphs may represent diverging lineages that evolved different radula morphologies in adaptation to alternative foraging substrates (Figure 2a). Further, the continuous secretion of the radula, which consists of numerous rows of chitinous teeth (Suppl. figure 1)^36^, enables drastic phenotypic plasticity in some snails^37,38^, showing that tooth shapes can be altered during radula secretion. However, such phenotypic plasticity does not seem to occur in *T. sarasinorum*. In fact, both ecomorphs can be found across both substrates, yet changes in radula morphology across teeth rows have never been observed in ~500 specimens (Suppl. table 1).

**Fig. 2:**
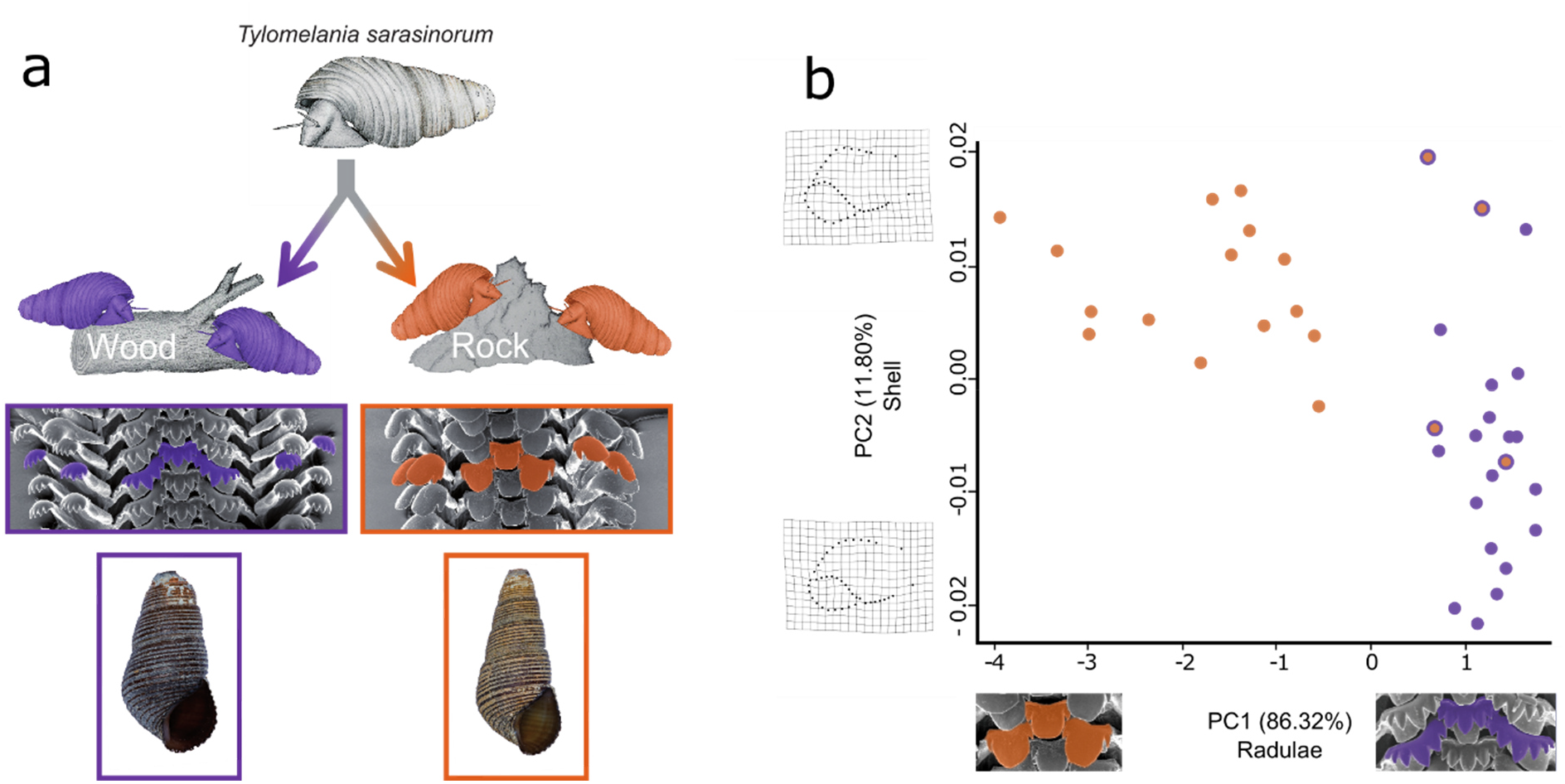
Habitat-correlated radula polymorphism in *T. sarasinorum*. a) Hypothesis: Radulae of *T. sarasinorum* evolved in adaptation to different microhabitats, i.e. submerged logs and rocks giving rise to diverging ecomorphs. Subtle differences in shell shape and coloration likely also exist. b) Scatterplot based on the two principal components of shell and radula shape that differ significantly between wood (purple) and rock (orange) ecomorphs. Thin plate splines visualize shape change explained by PC2 of shell shape. The center of each dot indicates the habitat from which individuals were collected while the outer ring indicates to which ecomorph it was assigned based on SEM inspection of the radula. Individuals of both ecomorphs are associated with alternative substrates and clearly differ in radula tooth morphology. However, less pronounced, more gradual differences in shell shape also exist.

Here we combine morphological analyses of the radula and the shell with tissue-specific transcriptomes to measure morphological and genetic divergence of sympatric *T. sarasinorum* ecomorphs. Our results indicate evolutionary divergence of ecomorphs. Divergence is most pronounced between radula transcriptomes, adding support to the hypothesis that the radula acts as key adaptive trait in *Tylomelania* radiations. Putative homologs of candidate genes for radula diversification also contributed to morphological diversification in vertebrate radiations. Our findings indicate that adaptive diversification can leave tissue-specific footprints of transcriptomic divergence, while morphological diversification in adaptive radiations may preferentially be achieved via a limited set of hotspot genes^39,40^ within conserved signaling pathways.

## Results and discussion

### Geometric morphometrics corroborates a habitat-correlated radula polymorphism

Although a habitat-correlated radula polymorphism of *T. sarasinorum* together with slight differences in shell morphology and coloration (Figure 2a) have previously been reported^41^, these patterns have not been systematically analyzed so far. Hence, we investigated whether radula morphs are indeed morphologically distinct and whether frequencies of ecomorphs are habitat-correlated at Loeha Island. To this end, we quantified variation in radula and shell morphology of 19 and 18 specimens collected on wood and rock substrates, respectively (See suppl. figure 2 for an overview of radula morphologies). We found that the frequency of both radula morphs differs significantly depending on the substrate from which they were collected (15/19 = 79% of specimens on rock are rock ecomorph; 18/18 = 100% of specimens on wood are wood ecomorph; p = 1.01*10^−6^; χ^2^ test). Furthermore, the first principal component (PC1) of radula shape, which accounts for 93.4% of the variance within the dataset, clearly separates *T. sarasinorum* ecomorphs (p < 0.001). No overlap in PC1 exists between radula morphs, even when individuals collected on opposing substrates where included in the analysis (Figure 2b). In contrast, differences in shell shape are both more subtle and more gradual. Although significant differences in shell shape exist in PC2 (p < 0.001), this axis summarizes a relatively small proportion of shell shape variance (11.8%) (Suppl. tables 2, 3). Furthermore, shell morphospaces of both ecomorphs are largely overlapping along this axis (Figure 2b). Taken together our data support a habitat-correlated radula polymorphism in *T. sarasinorum*. Consequently, *T. sarasinorum* ecomorphs represent a promising model to study the molecular basis of radula disparity and its role for lineage divergence in adaptive radiations of *Tylomelania*.

### Transcriptome sequencing and assembly

To gain insight into transcriptomic divergence of sympatric *T. sarasinorum* ecomorphs, we pooled tissues from four to five individuals and conducted RNAseq on four biological replicates of each mantle, radula formative tissue (Suppl. figure 1) and foot tissue from both ecomorphs. A single transcriptome was assembled from combined data of both ecomorphs. Removal of genes with low expression and clustering of sequences with high sequence similarity (>97%) reduced the number of contigs with gene status (assigned by Trinity) from 478,661 to 156,685 and increased N50 from 613 bp to 1229 bp (Table 1). Importantly, filtering did not affect assembly completeness (89%) based on a search for 843 conserved metazoan single-copy orthologs using BUSCO^42^, but decreased the rate of duplicated single copy orthologs from 9.4% to 7.5% (Table 1). Since N50, completeness and duplication rate are well within the range of recently published mollusk transcriptomes^43,44^, this assembly was used to analyze transcriptomic divergence between ecomorphs of *T. sarasinorum*. Only three out of four biological replicates were used in the subsequent analyses after pool1 of either ecomorph was identified as outlier in initial gene expression analyses (Suppl. figure 3)^45^.

**Table 1:** Assembly statistics of the raw and filtered assembly

|  | Trinity<br>genes | GC<br>(in %) | 'gene'<br>N50 | Complete<sup>a</sup><br>(in %) | Duplicated<sup>a</sup><br>(in %) |
| --- | --- | --- | --- | --- | --- |
| **Raw**<br>**assembly** | 478 661 | 45.2 | 613 | 89 | 9.4 |
| **Filtered**<br>**assembly** | 156 685 | 44.9 | 1 229 | 89 | 7.5 |

<sup>a</sup> According to BUSCO

### Transcriptome wide SNP data indicates divergence of *T. sarasinorum* ecomorphs

Lineage diversification in adaptive radiations of *Tylomelania* was hypothesized to be promoted by radula diversification in adaptation to alternative substrates^31,32^. Hence, we investigated whether sympatric radula morphs of *T. sarasinorum* with different habitat preferences represent diverging evolutionary lineages. To this end, population genetic analyses were carried out with PoPoolation2^46^ using a uniform coverage of 20x and 10% minor allele frequency (MAF) for SNP detection. In a total of 39,631,840 bases that passed the filtering steps, 517,825 putative SNPs could be identified. Of these putative SNPs, 6,366 SNPs (1.2%) in 2,572 transcripts (7.8% of all transcripts with putative SNPs) qualified as alternatively fixed between ecomorphs (F_st_ = 0 in all within-morph comparisons and F_st_ = 1 in all across-morph comparisons). Figure 3a depicts pairwise genetic differentiation between all pools, while Figure 3b shows the SNP-wise F_st_ distribution between both ecomorphs. Although the majority of genetic variation is shared between populations of both radula morphs at Loeha Island (median F_st_ = 0.14; mean F_st_ = 0.23), we observed an excess of highly differentiated loci and consistently higher F_st_ between pools of different ecomorphs (Figure 3). While median F_st_ in pairwise comparisons among pools of similar ecomorphs ranged from 0.016 to 0.048, it ranged from 0.143 to 0.188 among pools of different ecomorphs. Accordingly, our transcriptome-wide SNP data indicate evolutionary divergence of sympatric radula morphs of *T. sarasinorum* at Loeha Island. Although the F_st_ distribution indicates high differentiation of a few genomic regions in a background of shared genetic variation, pooled transcriptomic data does not allow to reliably distinguish between potential scenarios that may have given rise to this pattern. One possibility that could give rise to such patterns is divergence with gene flow^47,48^. During divergence with gene flow, a few loci under selection become fixed whereas genomic variation at sites of the genome that are not in strong linkage with selected loci are homogenized by gene flow, as long as reproductive isolation remains incomplete^47,48^. In fact, individuals with intermediate phenotypes and non-resolving phylogenies from mitochondrial markers indicate that gene flow may not only exist between ecomorphs, but even among *T. sarasinorum* and other species^34,49^. However, other scenarios like divergence without gene flow combined with selective sweeps, potentially following secondary contact, may result in similar patterns, albeit with increased absolute divergence in regions that are not linked to outlier loci. Genomic data comprising individuals from other sites and ideally other species would be required to investigate population history and gene-flow among divergent lineages to decide between alternative explanations.

**Fig. 3:**
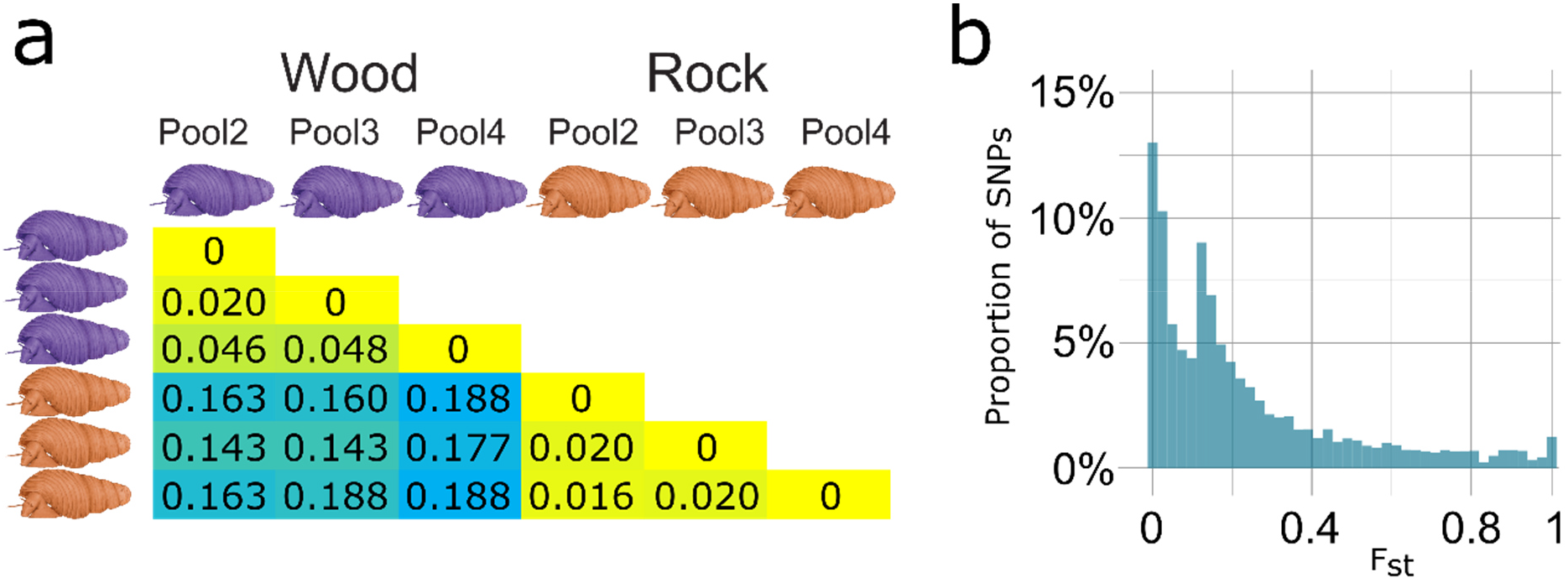
Divergence of *Tylomelania sarasinorum* ecomorphs. a) For all pairwise comparisons of pools from wood (violet) and rock (orange), differentiation measured as median SNP-wise Fst, is depicted. The degree of differentiation from low to high is indicated by color change from yellow to blue. b) Distribution of SNP wise Fst between different ecomorphs for 517,852 putative SNPs (60x (20x per pool) minimum coverage, 10% MAF, all pools of each ecomorph combined). While some SNPs exhibit high differentiation, the majority of variation is shared among both ecomorphs.

### Ecomorphs differ in gene expression across all investigated tissues

Regulatory evolution resulting in divergent gene expression plays a key role in adaptation and speciation^29,50^. Although gene expression is known to be highly tissue dependent, our knowledge of tissue-specific transcriptomic divergence is still in its infancy^28,51^. To shed light on gene expression divergence between *T. sarasinorum* ecomorphs, gene expression of foot tissue, shell forming mantle and radula forming tissue of both ecomorphs was analyzed using the pipeline included in Trinity v2.1.1^52,53^.

In accordance with previous work, foot and mantle form sister clusters to the exclusion of the radula cluster and biological replicates cluster tightly together in both PCA and hierarchical clustering, i.e., without a priori assumptions concerning group affiliation (Figure 4a,b)^45^. Within tissues, samples of different ecomorphs form separate clusters, indicating divergence in gene expression across all investigated tissues (Figure 4a, b). Finally, we analyzed genes that are highly differentially expressed between identical tissues of both ecomorphs (false discovery rate (FDR) ≤ 10^−10^; fold change (FC) ≥ 4; Figure 4c). Although overall fewer genes are expressed in the radula than in the other two tissues, more genes are highly differentially expressed between the radula forming tissues (n = 536, 0.81% of genes that are expressed in at least one radula tissue) than between mantle (n = 436, 0.34%) or foot tissues (n = 424, 0.42%) of the two ecomorphs. Stronger morphological disparity in the radula compared to the two other tissues is thus mirrored by more pronounced differences in gene expression, which indicates that regulatory evolution contributes to morphological radula disparity.

**Fig. 4:**
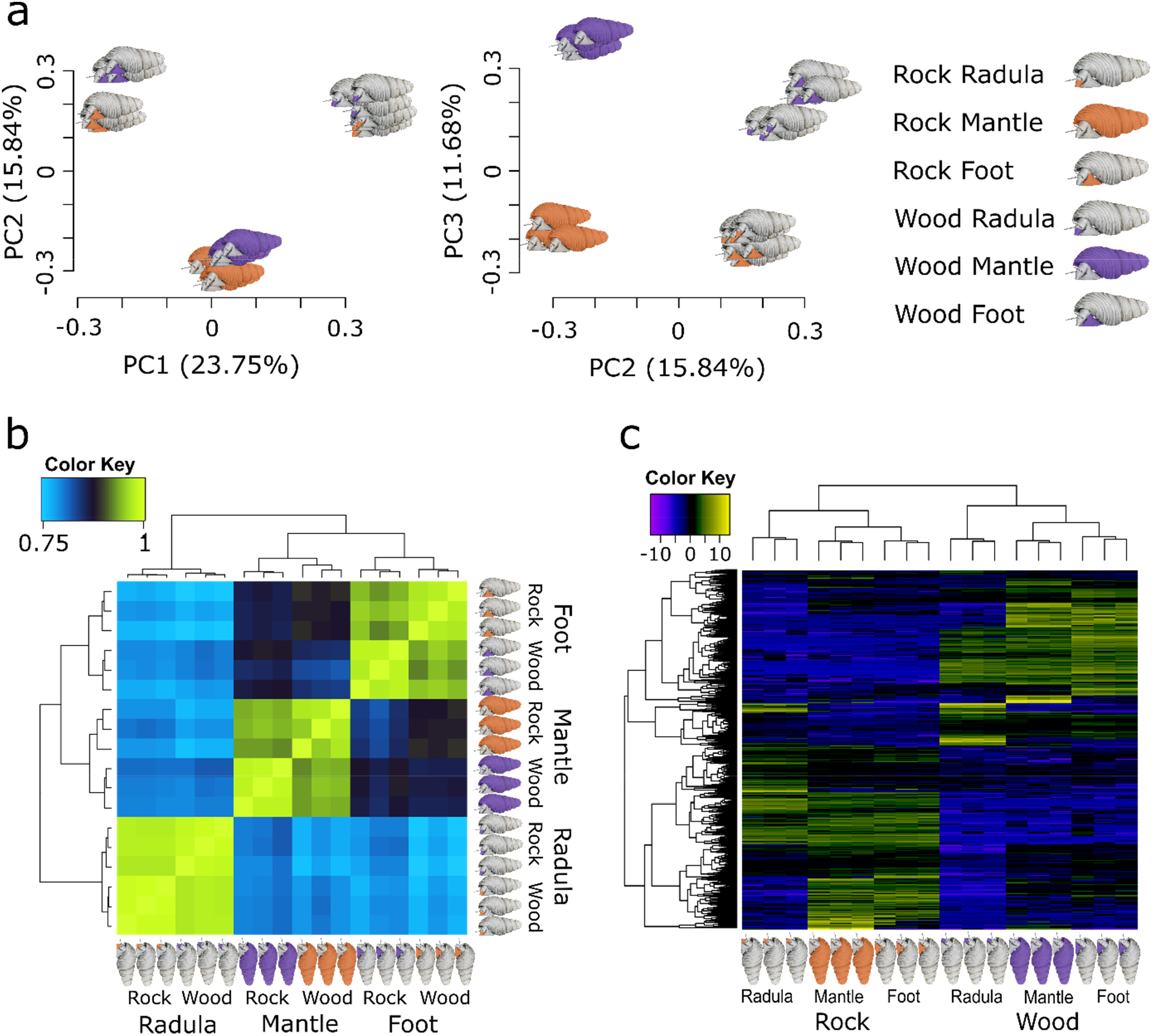
Divergence of gene expression between *T. sarasinorum* ecomorphs. a) depicts a principal component analysis (PCA) of gene expression in radula, mantle and foot tissue from wood (violet) and rock (orange) ecomorphs. The first and second principal components (PCs) primarily separate different tissue types, while the third PC separates tissues derived from different ecomorphs. b) Hierarchically clustered Spearman correlation matrix of gene expression (log_2_ transformed CPM). Samples with more similar gene expression cluster together in the matrix and the hierarchical clustering tree (left and top) and are colored yellow in the heatmap. c) shows differentially expressed genes between identical tissues of both ecomorphs. Genes are displayed as horizontal lines across samples (columns) in a heatmap of hierarchically clustered, highly differentially expressed (p ≤ 10^−10^, FC ≥ 4) genes between identical tissues of wood (violet) and rock (orange). Genes with similar expression across samples cluster together in the hierarchical clustering tree on the left, while samples with similar gene expression cluster together in the clustering tree on the top. Overexpressed genes in a sample are colored yellow in the heatmap, while underexpressed genes are displayed in blue.

### Elevated divergence of radula transcriptomes supports the radula’s role as key adaptive trait

Selection experiments have shown that strong selection can result in rapid tissue-specific transcriptomic divergence^54^. We thus hypothesized that if diversifying selection on the radula drove divergence of *T. sarasinorum* ecomorphs, divergence of the radula transcriptome would be stronger than transcriptomic divergence of other tissues. To further investigate the contribution of changes in gene regulation and protein coding sequences to radula evolution, we determined tissue-wise transcriptomic divergence in both gene expression and coding sequences.

Divergence in gene expression was measured as the proportion of highly differentially expressed genes (FDR ≤ 10^−10^) between the same tissue types of both ecomorphs. In this context, genes which are universally differentially expressed across all sampled tissues are uninformative and were excluded from the analysis. We found that divergence in gene expression is significantly higher in the radula than in the mantle (97% higher; p < 10^−5^; Fisher’s exact test) and in foot (85% higher; p < 10^−5^), while no significant difference exists between the latter two (6% higher in foot, p = 0.42) (Figure 5a). The radula also has the highest proportion of highly differentially expressed genes at lower false discovery rates (e.g. FDR ≤ 10^−100^). Only when the criteria for DE genes are relaxed and higher false discovery rates are accepted, foot gains the highest rate of differentially expressed genes (FDR ≤ 10^−5^: radula vs. mantle: 3,5% higher in the radula, p = 0.42; foot vs radula: 15,5% higher in foot, p < 10^−5^; foot vs mantle: 19,5% higher in foot, p < 10^−5^).

**Fig. 5:**
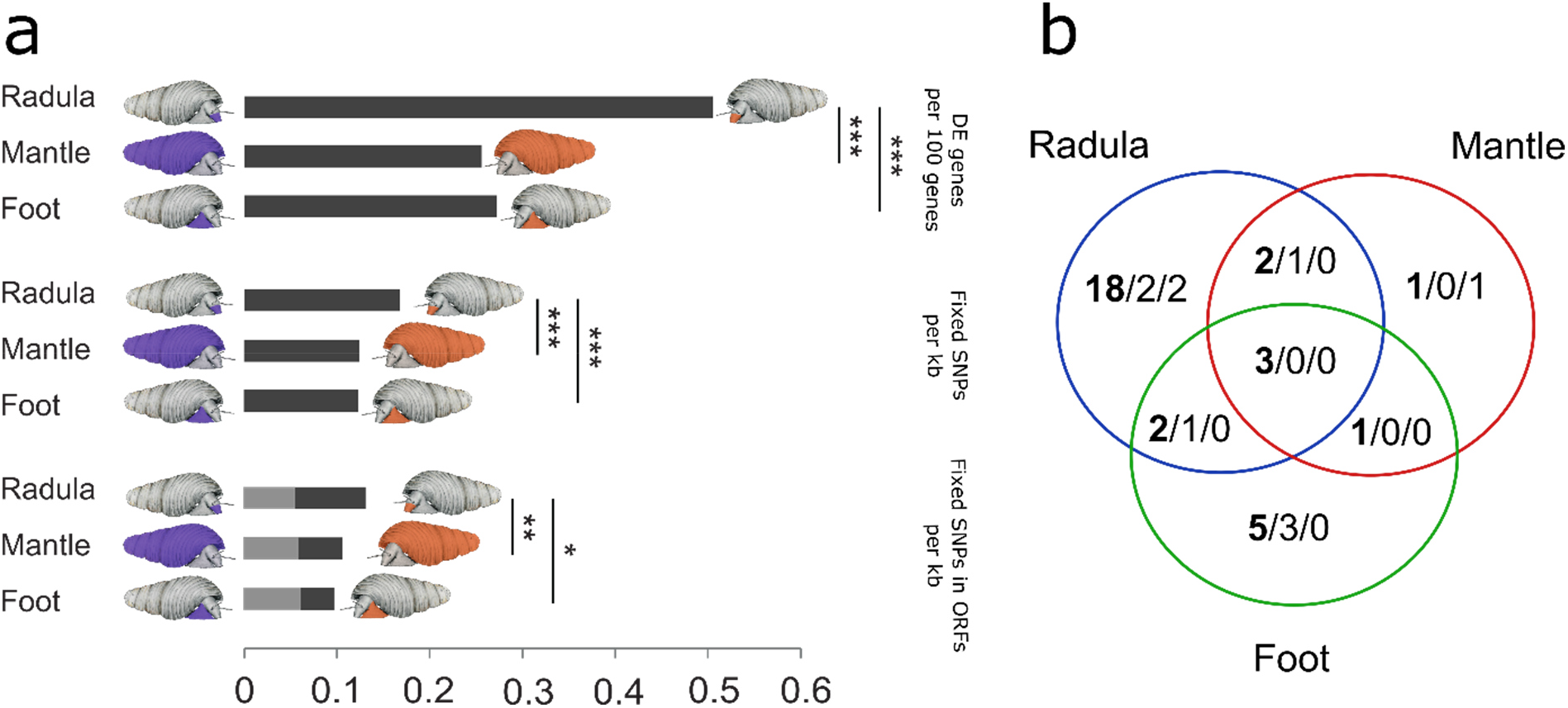
Tissue-wise transcriptomic divergence of *Tylomelania sarasinorum* ecomorphs. The proportion of highly differentially expressed (DE) genes between rock and wood ecomorphs, frequency of alternatively fixed SNPs, and frequency of such SNPs in ORFs (black = synonymous; grey = non-synonymous) is shown for genes expressed in each tissue separately. For the proportion of alternatively fixed SNPs, only genes that are expressed (FPKM ≥ 1) in the respective tissue, but are not expressed across all tissues are included. Similarly, only genes that are not differentially expressed between ecomorphs across all tissues were considered for tissue-wise rates of differentially expressed genes. Significant differences between tissues are indicated by asterisks (* = p ≤ 0.05; ** = p ≤ 0.01; *** = p ≤ 0.001). Divergence in both gene sequences and gene expression is significantly higher in genes that are expressed in the radula. b) Venn graph illustrating the position of alternatively fixed SNPs in genes that are also highly differentially expressed between at least one pair of identical tissues of both ecomorphs. The total number of SNPs in highly DE genes is shown first and in bold, followed by the number of synonymous and non-synonymous SNPs in these genes. The majority of alternatively fixed SNPs lie outside of ORFs and are found in genes that are only highly differentially expressed between radula forming tissues.

Similar to the proportion of differentially expressed genes, the frequency of alternatively fixed SNPs in transcripts of genes that were not expressed across all tissues (FPKM < 1, i.e. less than one mapped fragment per kilobase of transcript per million mapped reads, in all biological replicates of one tissue) is significantly higher in the radula than in mantle (~ 34.4% higher in radula, p < 10^−5^) or foot (36.6% higher in radula, p < 10^−5^), but does not differ significantly between the latter two (1.6% higher in mantle, p = 0.84) (Figure 5a). This pattern remains unchanged when only alternatively fixed SNPs within open reading frames (ORFs) are considered (radula vs mantle: 23.4% higher in radula, p = 0.036; radula vs foot: 34.7% higher in radula, p = 0.005; mantle vs foot: 9.2% higher in mantle, p = 0.30). No significant differences among tissues were found when the analysis was restricted to non-synonymous, alternatively fixed SNPs (radula vs. mantle: 6.5% higher in mantle, p = 0.72; radula vs foot: 11.3% higher in foot, p = 0.51; mantle vs foot: 4.5% higher in foot, p = 0.65) (Figure 5a). Finally, when both datasets were combined, the majority of genes with alternatively fixed SNPs that were also highly differentially expressed between ecomorphs were only highly differentially expressed between the radula tissues (Figure 5b, but see suppl. figure 4 for lower DE threshold).

Our findings indicate that both the rate of highly differentially expressed genes and the frequency of alternatively fixed SNPs are significantly higher in genes expressed in the radula, compared to genes expressed in the other investigated tissues, which is in accordance with stronger diversifying selection on the radula transcriptome. An alternative explanation for a higher rate of differentially expressed genes in the radula would be that the possibility for more precise sampling of radula forming tissue reduces noise in gene expression data, which favors the detection of differentially expressed genes. However, higher sequence divergence of the radula transcriptomes cannot be explained in a similar fashion. Hence, tissue-specific patterns of transcriptomic divergence add support to the hypothesis that diversifying selection on the radula in the course of adaptation to alternative substrates promotes lineage diversification in the adaptive radiations of *Tylomelania*^31^.

Although genes expressed in the radula exhibited significantly higher sequence divergence compared to mantle and foot, no significant differences were detected for non-synonymous mutations (Figure 5a). Sequence divergence outside of ORFs may reflect divergent non-coding RNAs or untranslated regions of protein coding transcripts, both of which can be linked to post-transcriptional regulation^55,56^. Overall, regulatory evolution appears to dominate divergence of *Tylomelania* ecomorphs as indicated by highly divergent gene expression across all tissues, which is most pronounced between radulae and higher divergence in untranslated than translated regions, both in general and in transcripts of highly DE genes (Figure 5a,b). These findings are in accordance with the expectation that the relative contribution of protein coding evolution to phenotypic disparity ceases over time, because selection favors regulatory change that can avoid deleterious pleiotropic effects^39,40^. Additionally, our results suggest that divergence of ecomorphs is likely polygenic, which is in line with results from other study systems. For example, regulatory evolution contributed to ecological divergence in East African cichlids, Darwin’s finches and sticklebacks ^2,18,21,57^, and polygenic selection gave rise to convergent gene expression in lake whitefish radiations in Europe and North America^29^.

### Functional enrichment of differentially expressed genes hints at tetrapyrrole synthesis underlying shell color disparity

To gain insight into dominant molecular functions (MF), cellular components (CC) and biological processes (BP) in genes contributing to divergence between ecomorphs, transcripts were functionally annotated with the Trinotate annotation pipeline (https://trinotate.github.io/) and gene ontology (GO) enrichment analyses were carried out with GOseq^58^. Similar to previously published mollusk transcriptomes^43,45^, only a minority of transcripts could be annotated (n = 29.139; 19 %) and GO terms were assigned to 13% (n = 20,864) of all sequences in the final assembly. The only enriched GOs among all transcripts with alternatively fixed SNPs were the BP term “biological process” and the CC term “cellular component”, indicating that non-synonymous SNPs accumulated in transcripts with unknown functions. In contrast, highly differentially expressed genes between identical tissues of the two ecomorphs were enriched in multiple MF (n = 9) including “carbohydrate binding”, “heme binding” and “tetrapyrrole binding” (Suppl. table 5). The enriched GO-term “carbohydrate binding” is unsurprising within this context, because the radula is primarily made up of the polysaccharide chitin and proteoglycans are important constituents of the molluscan shell^59^. In contrast, implications of the enriched MF “heme binding” and “tetrapyrrole binding” are not as obvious. Interestingly, tetrapyrroles are important molluscan shell pigments^60,61^ and synthesized in the heme pathway^62^. Tetrapyrrole binding genes that were differentially expressed between ecomorphs are primarily expressed in the shell building mantle and foot of the rock morph and mostly differentially expressed between the mantle tissues of both ecomorphs (Suppl. figure 5). Although further investigations targeting color differences between *T. sarasinorum* ecomorphs are needed, enrichment of these functions suggests that differential gene expression in the tetrapyrrole synthesis pathway in the mantle may underlie shell color differences between the ecomorphs.

### Candidate genes for radula disparity include cell-cell signaling genes involved in craniofacial diversification in vertebrate radiations

To investigate individual genes that contributed to radula diversification, two non-overlapping sets of candidate genes were generated based on i) differential expression and ii) non-synonymous protein coding sequence divergence. Genes that were highly differentially expressed between the radulae of the two ecomorphs (FDR ≤ 10^−10^; FC ≥ 4), but not differentially expressed (FDR ≥ 10^−5^; FC ≤ 4) between mantles or foot tissues, were chosen as expression-based candidate genes (n = 230). The second set of candidate genes (n = 538) was composed of genes that were expressed in the radula of both ecomorphs and carried alternatively fixed non-synonymous SNPs. To further narrow down the list of candidates, we focused on genes involved in gene regulation and cell-cell signaling, because both regulatory as well as protein coding evolution of these genes may determine when and where radula tooth matrix is secreted.

While most genes with alternatively fixed non-synonymous SNPs only had one such SNP (66%), a maximum of 12 such SNPs (and 10 synonymous) was found in Rho GTPase activating protein 21 (*rhg21*) (Suppl. figure 6). Rho GTPase activating proteins are important activators of Rho family GTPase signaling. Rho family GTPase signaling interacts with notch signaling and regulates various cellular functions, including cytoskeletal reorganization in response to extracellular stimuli^63–66^. Coordinated reorganization of the cytoskeleton is particularly interesting with respect to the radula polymorphism of *T. sarasinorum*, because odontoblasts undergo pronounced shape changes during radula tooth secretion, and modification of their cell shape likely influences tooth morphology^36^. In addition to changing odontoblast cell shapes, modified cytoskeletons may change the localization of chitin synthesis via altered actin filament guidance of a lophotrochozoan-specific chitin synthase with a myosin head^67^ that is expressed in radula forming tissue^45^. The number of non-synonymous alternatively fixed SNPs per kb of ORF in *rhg21* corresponds to a 32.2-fold and 2.37-fold increase in frequency of such SNPs compared to the average of all transcripts in the analysis and all transcripts with alternatively fixed SNPs, respectively. Unless mutation rates of genes like *rhg21* are substantially increased, their alleles diverged before the divergence of *T. sarasinorum* ecomorphs and either persisted as standing genetic variation, or were introgressed from different lineages. Gene flow among diverging lineages and even introgression from more distantly related species is common in adaptive radiations and may generate and maintain genetic variation at loci underlying adaptive traits^2,18,20,21,23,24,68–72^. Since previous studies indicate abundant hybridization among species of *Tylomelania*^31,34^, extraordinary divergence of a few genes like *rhg21* between *T. sarasinorum* ecomorphs (Suppl. figure 6) indicates that selection on highly divergent introgressed alleles may also contribute to lineage divergence in adaptive radiations of *Tylomelania*. Genomic data from across the radiation could be used to test this hypothesis, which, if confirmed, would add further support to a combinatorial view on speciation and adaptive radiation (reviewed in ^73^).

SNP-based candidates further included a transcript that was annotated as *neurogenic notch locus 1* (*notch1*) and a transcript encoding strawberry notch (1 non-syn; 5 syn), whose role in the notch signaling pathway is still unclear^74^. A putative homolog of *notch1* was also found among the expression-based candidate genes, together with the morphogen *hedgehog* (*hh*). Both the notch and the hedgehog signaling pathway are conserved across the bilaterians and interact during developmental tissue patterning^75,76^. Interestingly, the hedgehog signaling pathway mediates both fixed as well as phenotypically plastic effects on jaw morphology in East African cichlids^77–79^. Hedgehog (*hh*) further regulates bone morphogenetic protein (BMP) expression in several metazoan lineages^76,80,81^ and regulatory evolution resulting in divergent expression of BMPs played a pivotal role for craniofacial diversity in both Darwin’s finches and East African cichlids^26,27,82,83^. In *T. sarasinorum, hedgehog* (*hh*) is overexpressed in the radula of the rock morph and a BMP that is most similar to *gbb*/*BMP5-8* is only expressed (FPKM ≥ 1) in the radula of the rock morph. The only homeobox gene found among candidate genes was annotated as *aristaless related homeobox protein* (*arx*) (4 non-syn; 2 syn). In our dataset, a*rx* is only expressed in the radula tissue and carries four non-synonymous alternatively fixed SNPs. Similar to notch signaling, we previously found that *arx* likely plays an important role for radula formation^45^. Interestingly, two non-synonymous substitutions in the *aristaless-like homeobox 1 transcription factor* (*ALX1*), which has an important role for craniofacial development in vertebrates, promoted beak diversification in Darwin’s finches^21^.

In summary, we find that several close relatives and putative homologs of genes that contributed to diversification of beaks in Darwin’s finches and/or yaws of East African cichlids might also be involved in the adaptive diversification of the radula. Although, given similar gene regulatory networks, evolutionarily relevant mutations are expected to accumulate in so-called hotspot genes^39,40^, the radula does not share the developmental basis that jaws and beaks have in common^84^. Nonetheless, our observations might be explained by a relatively restricted and highly conserved set of tissue patterning cell-cell signaling pathways^85^ that contain a limited set of genes that have the potential to rapidly generate potentially adaptive morphological diversity without fatal pleiotropic effects^39,40,76,86^. While a large number of candidate genes in this study calls for further verification, our results indicate that diversification of foraging organs in adaptive radiations might be achieved via a limited set of cell-cell signaling genes that are particularly prone to rapid adaptive diversification.

## Conclusions

This study confirms habitat-correlated radula disparity in *T. sarasinorum*, shows evolutionary divergence of ecomorphs and corroborates the hypothesis that adaptive diversification of the radula drives lineage divergence in adaptive radiations of *Tylomelania*. Exceptional sequence divergence of some genes may be a sign of older, potentially introgressed, variation, but needs to be further investigated using population genomic data from other sites and including related species. More generally, our findings shed light on tissue-wise transcriptomic divergence and indicate that adaptive diversification can leave tissue-specific footprints. Finally, overlapping gene sets appear to underlie rapid adaptive diversification of foraging organs in radiations of fishes, birds and snails, which is an aspect that requires further investigation in the future to get a better understanding of the genetic mechanisms generating functional diversity within adaptive radiations.

## Materials and Methods

### Specimen and tissue collection

Adult specimens of *Tylomelania sarasinorum* were collected from submerged wood and rock substrates at the northern shore of Loeha Island (Lake Towuti, South Sulawesi, Indonesia; 2.76075 S 121.5586 E). All snails were collected in close proximity to each other and kept in buckets filled with lake water for a few hours before they were dissected in the field. Tissue samples of radula forming tissue, mantle edge, and foot muscle were directly stored in RNAlater to ensure RNA preservation. Before any samples were pooled for RNA extractions, radula forming tissue was separated from the rest of the radula (Suppl. figure 1) and radula morphs of all individuals were inspected using scanning electron microscopy.

### Morphological analyses

Shell shape and radula meristic were assessed for a total of 37 adult specimens from the collection of the Natural History Museum Berlin (Suppl. figures 1, 7). Specimens were chosen from lots that had been sampled randomly by hand from wooden (n = 19) and rocky substrate (n = 18).

Variation in shell shape was quantified using landmark based geometric morphometrics methods. To this end, specimens were placed on sand-filled trays and photographed with the aperture facing upwards using a SatScan collection scanner (SmartDrive Limited). Eight landmarks were placed on the whorls and aperture (see Suppl. figure 7a for details). Round structures of the aperture and the first and second whorl were outlined by four sliding semilandmarks (Suppl. figure 7a). Landmarks were placed using the software tpsDIG2^87^. Differences in size and rotation were removed from the data with a Procrustes superimposition (gpagen function in the geomorph package^88^). A principal component analysis (PCA) was calculated on the Procrustes residuals using the plotTangentSpace function (r package geomorph^88^). T-tests were calculated for principle components that explained more than 5% of the total variance, to test for significant differences in shell shape between ecomorphs. Radulae were dissected from the headfoot of the animals and surrounding tissue was digested with 500µl lysis buffer^89^ and 10µl proteinase K at 55°C overnight. Afterwards, radulae were cleaned with ethanol and treated for 2 seconds in an ultrasound bath. Radulae were mounted on electron microscope stubs and sputter coating was carried out with the Quorum Q150RS Sputter Coater using the manufacturer’s program number 2.

The number of teeth was counted and maximum width and total height of the central denticle as well as the total width of the rachis base were measured with the software ImageJ^90^ (Suppl. figure 7b). Subsequently, ratios of central denticle width/total height and rachis width were calculated. A PCA was carried out with these ratios and the number of denticles of the rachis (the central tooth). Two tailed t-tests were used to evaluate morphological differences between ecomorphs for each PC that explained more than 5% of the total variance.

### Sample preparation and sequencing

Nineteen individuals of the *T. sarasinorum* wood morph were grouped into three pools of five and one pool of four individuals (data already used in^45^), and 20 individuals of the *T. sarasinorum* rock morph were grouped into four pools of five individuals. Tissue samples of individuals in each pool were weighed (Mettler AT 261 scale), and similar amounts of each individual were pooled, resulting in four biological replicates of each tissue. Tissue was homogenized with a Precellys Minilys, and total RNA was extracted using two alternative protocols. Since larger amounts of foot tissue were available, RNA was extracted from foot muscle with a TRIzol® extraction according to the manufacturer’s protocol. However, to extract RNA from minute amounts of radula formative tissue and mantle edge, a customized protocol of the RNeasy Plus Micro Kit (Qiagen) was employed^45^. Briefly, remaining tissue fragments were digested with proteinase K following mechanical homogenization. Subsequently, lysis buffer was added to allow efficient DNA removal with gDNA spin columns. Amount and quality of extracted total RNA was inspected using Agilent’s 2100 Bioanalyzer. *Tylomelania sarasinorum* rRNA carries a “hidden break”, which means that the 28S rRNA easily disintegrates into two smaller fragments. This led to a sharp 18S band, but a much reduced or lacking 28S rRNA peak in our samples. Hence, RNA integrity (RIN) estimates were not applicable. Nonetheless, samples showed no signs of degradation or DNA contamination. Messenger RNA was enriched with poly (A) capture using NEXTflex™ Poly (A) Beads, and strand-specific libraries were built using the NEXTflex™ Rapid Illumina Directional RNA-Seq Library Prep Kit (Bioo Scientific) with modifications suggested by Sultan et al. (2012). Quality and concentrations of libraries were evaluated using Agilent’s 2100 Bioanalyzer and qPCR (Kapa qPCR High Sensitivity Kit). Libraries had average fragment sizes between 350-500 bp and were sequenced (150 bp, paired end) on an Illumina NextSeq sequencing platform at the Berlin Center for Genomics in Biodiversity research (BeGenDiv).

### Transcriptome assembly

Raw sequences were trimmed with a quality threshold of 30, minimum read length of 25 bp, and all Ns were removed using sickle^92^. Adapter sequences were subsequently removed with cutadapt^93^, which generated a final dataset consisting of 941 million paired end reads (Suppl. table 5). Trinity v2.1.1^52,53^ was run in strand-specific mode with a minimal transcript length of 250 bp, in silico read normalization (max. read coverage = 50), and two-fold minimal kmer coverage to generate a single assembly of all tissues of both ecomorphs. Quality-filtered adapter trimmed reads of each sample were mapped to the transcriptome using bowtie2^94^, followed by abundance estimation with RSEM^95^. Since abundance of rRNA mostly reflects polyA capture success, ribosomal RNA (rRNA) was removed following identification with a BLAST search using 28S rRNA (*Brotia pagodula*; HM229688.1) and 18S rRNA (*Stenomelania crenulata*; AB920318.1) as query sequences. Pool1 mantle and pool1 radula of both ecomorphs were removed from further analyses after they were identified as outliers in principal component analysis (PCA) of log_2_ transformed counts per million mapped reads (cpm) (Suppl. figure 3). The cause for this observation is likely a combination of lower yield of total RNA in the first extractions, which led to a decrease in library complexity, and deeper sequencing of pool1 (Suppl. table 5). A batch effect might also have contributed to this observation, because pool1 mantle and pool1 radula of both ecomorphs were sequenced separately from all other samples. The assembly was subsequently filtered by expression (FPKM ≥ 1, i.e. at least one mapped fragment per kilobase of transcript per million mapped reads), using a script provided in the Trinity pipeline. CD-HIT version 4.6^96^ was used to cluster the longest isoforms of all “trinity genes” based on sequence similarity (97% sequence identity threshold; 90% minimum alignment coverage of the shorter sequence), and the longest transcript of each cluster was retained. Quality filtered, adapter trimmed reads of both ecomorphs were re-mapped to the remaining transcripts, and transcripts with very low expression (FPKM ≤ 1) were removed to create a final assembly. BUSCO v1.1b1^42^, was employed to generate estimates of transcriptome completeness, redundancy, and fragmentation by searching for 843 known metazoan single copy orthologs. Since BUSCO indicated that transcriptome completeness was not negatively affected by the abovementioned filtering steps, the final assembly was chosen for further analyses.

### Gene expression analysis

Gene expression analysis was performed using the pipeline included in Trinity v2.1.1^52,53^. Briefly, quality-filtered adapter-trimmed reads of each sample were mapped to the final assembly using bowtie2^94^, followed by abundance estimation with RSEM^95^. Differentially expressed genes (FDR ≤ 10^−5^; FC ≥ 4) and highly differentially expressed genes (FDR ≤ 10^−10^; FC ≥ 4) were determined for all pairwise morph and tissue comparisons using edgeR^97^.

### Annotation

Transcripts in the final assembly were functionally annotated using the Trinotate annotation pipeline (v3.0.1). Results were imported into the Trinotate-SQLite database, and the annotation report was generated using default parameters. The identities of *T. sarasinorum* genes that are mentioned by name in this manuscript were further varified by searching proteins matching *T. sarasinorum* open reading frames in the UniProt database using BLASTX and manually inspecting alignments of the 10 best hits with an E-value of 10^−10^ or lower for which the alignment covered at least 60% of the database sequence.

### Ecomorph divergence

PoPoolation2^46^ was used to gain insight into divergence of *T. sarasinorum* ecomorphs. Duplicate reads, reads that did not map as proper pairs, and low quality alignments (mapping quality < 20) were removed from mappings using SAMtools v1.3^98^ and Picard Tools (http://broadinstitute.github.io/picard/). Subsequently, mappings of different tissues of the same pool (same individuals) were merged. To reduce biases in SNP detection caused by variance in gene expression, a uniform coverage of 20x for each pool was generated by subsampling mapped reads (without replacement) and removing all sites with a coverage <20x. SNPs were called at a minor allele frequency (MAF) of 10%, i.e. 12 calls in a total of 120 calls per site across all six pools. SNPs with lower MAF were discarded to remove potential sequencing errors and uninformative SNPs^99^, which increases the accuracy of allele frequency estimations^100^. SNP-wise F_st_ was calculated for all pairwise comparisons between pools using PoPoolation2^46^. Median pairwise F_st_ were estimated from all SNPs for each pairwise comparisons of pools. Median Fst and SNP wise F_st_ distributions between ecomorphs were calculated based on combined pool-wise allele counts for which a .sync file with combined allele counts for each the rock and the wood morph were generated. This resulted in a coverage of 60x (3 pools × 20x coverage per ecomorph). MAF was retained at 10%. Synonymous and non-synonymous mutations were determined using the *syn-nonsyn-at-position.pl* script included in PoPoolation v1.2.2 based on the longest ORF per gene and using a merged mapping file combining read mappings of all pools. Although PoPoolation is not recommended for processing pooled transcriptome data, because expression differences between individuals and alleles may introduce additional variation compared to sequencing pooled DNA (Pool-Seq)^46,100^, similar approaches have successfully been employed in numerous studies^54,101–104^. Additionally, high repeatability among biological replicates in this study supports the validity of our approach.

### Estimating tissue-specific transcriptomic divergence

To evaluate whether transcriptomic divergence between ecomorphs differed depending on the tissue, tissue-wise divergence in gene expression and coding sequences was determined. The proportion of genes that were highly differentially expressed between identical tissues of the ecomorphs (FDR ≤ 10^−10^; FC ≥ 4) was calculated for each tissue. Genes that were highly differentially expressed between ecomorphs across all tissues were excluded from the analyses. Likewise, the frequency of alternatively fixed SNPs was determined for genes expressed (FPKM ≥ 1) in each tissue, excluding genes that were expressed across all tissues. Genes were regarded as expressed in a certain tissue when they were expressed (FPKM ≥ 1) in at least one pool of that tissue. Differences in the proportion of differentially expressed genes and in the frequency of alternatively fixed SNPs between tissues were evaluated using Fisher’s exact test.

### Gene ontology enrichment

Gene ontology (GO) enrichment analyses were carried out to determine dominant functions of genes with alternatively fixed non-synonymous SNPs and of all genes that were highly differentially expressed (FDR ≤ 10^−10^; FC ≥ 4) between identical tissues of the two ecomorphs. For all transcripts in the final assembly, GO assignments and parental terms were extracted from the Trinotate annotation report using the script included in the Trinotate-2.0.2 pipeline. GOseq^58^ was used to identify enriched GOs in genes that were differentially expressed between the same tissues of different ecomorphs against a background of all genes in the final assembly. Additionally, enriched gene ontologies were identified in genes with alternatively fixed non-synonymous SNPs against a background of all genes that had bases that passed the filtering for coverage in the PoPoolation pipeline. Significantly enriched gene ontologies with a false discovery rate FDR ≤ 0.05 were summarized and redundant terms were removed (allowed similarity: 0.5) with REVIGO^105^.

### Identification of candidate genes

Alternatively fixed non-synonymous SNPs were used to identify candidate genes for adaptive divergence. Thresholds for outlier detection are always to some extent arbitrary and depend on the choice of MAF that is accepted as informative to detect patterns of selection^99,103^. In addition to demography and stronger purifying selection in the transcriptome resulting in different effective mutation rates^106^, core assumptions of models employed for pooled genomic data may be violated by a larger margin of error in allele frequency estimation from pooled RNA compared to pooled DNA due to variation in gene expression between individuals and even alleles^100^. Accordingly, previous studies based on pooled transcriptomic data mostly used quantile based approaches for outlier detection^101,103^. We used the most conservative approach available to us and only chose alternatively fixed SNPs, i.e. SNPs with F_st_ = 0 in all within-morph comparisons and F_st_ = 1 in all across-morph comparisons (98.8% percentile). Genes that carried non-synonymous alternatively fixed SNPs and were expressed in both radula forming tissues were determined as candidate genes for radula divergence. Finally, genes that were highly differentially expressed between the radulae of the two ecomorphs (FDR ≤ 10^−10^; FC ≥ 4), but not differentially expressed (FDR ≥ 10^−5^; FC ≤ 4) between mantles or foot tissues, were collected as candidate genes for radula shape divergence.

## Data availability

Sequence data and additional information are available at the NCBI Sequence Read Archive (SRP134819, ###) and BioProject (BioProject ID: PRJNA437798; BioSample accessions: SAMN08685289 - SAMN08685300 and SAMN13841508 - SAMN13841519).

## Acknowledgements

We thank Isabelle Waurick and the BeGenDiv for assistance in the lab, and all members of the Hofreiter Lab, von Rintelen Lab and David Garfield for helpful discussions. LIPI (Indonesian Institute of Sciences) and RISTEK (Indonesian State Ministry of Research and Technology) kindly issued the permits to conduct research in Indonesia. We would further like to thank Ristiyanti M. Marwoto (Museum Zoologicum Bogoriense, LIPI, Cibinong) for her continuous support of the project. This work was financed by the German Research Council (DFG) (grant number: Ri 1738/9-1). The authors declare no conflict of Interest.

