## Supplement for "Radula diversification promotes ecomorph divergence in an adaptive radiation of freshwater snails"

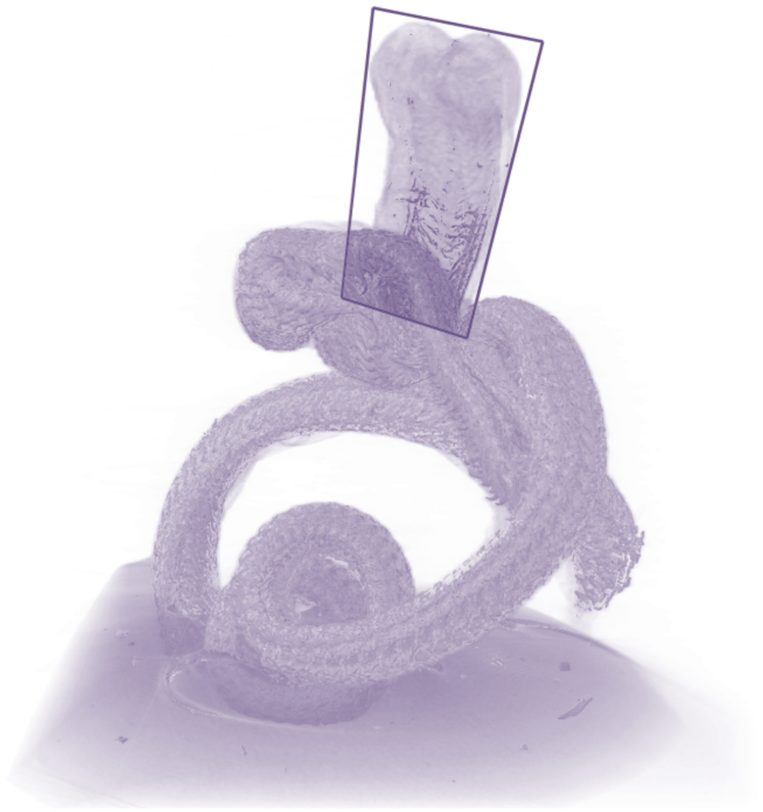

**Suppl. figure 1:** Micro-CT image of a dissected *Tylomelania* radula. The sampled radula forming tissue, where the radula is continuously secreted, is indicated by a box. Color differences represent differing densities and show starting tooth hardening (darker areas).

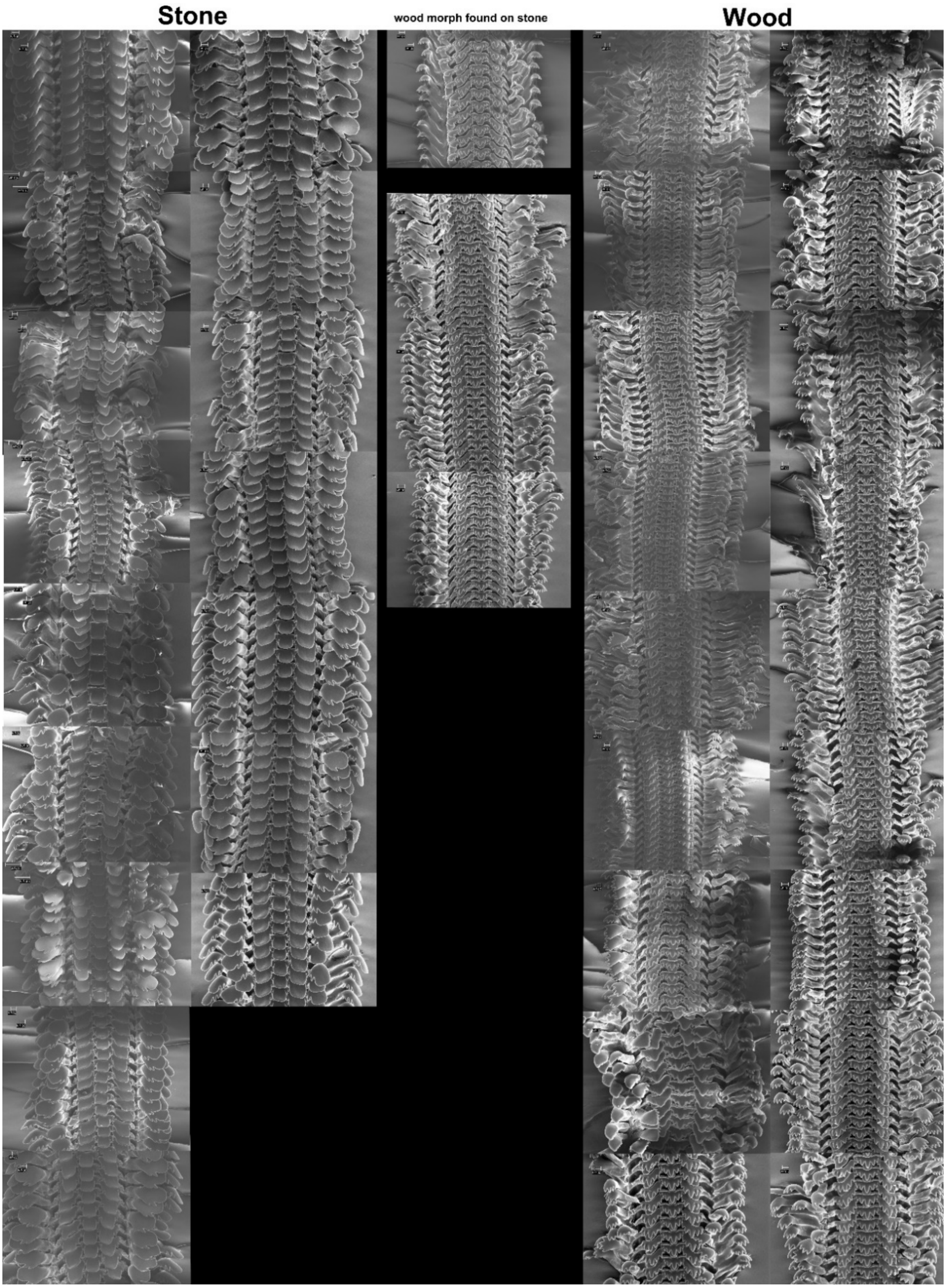


**Suppl. figure 2:** Overview of all radulae of individuals that were randomly collected from wood and rock substrates for morphological analyses. SEM images of radulae from *T. sarasinorum* rock morphs collected from rock (left), wood morph collected on wood (right) and wood morph collected on rock (middle) are shown. All individuals are shown, but one rock individual that was collected could not be used for further morphological analyses, because of insufficient resolution of morphological features required for measurements.

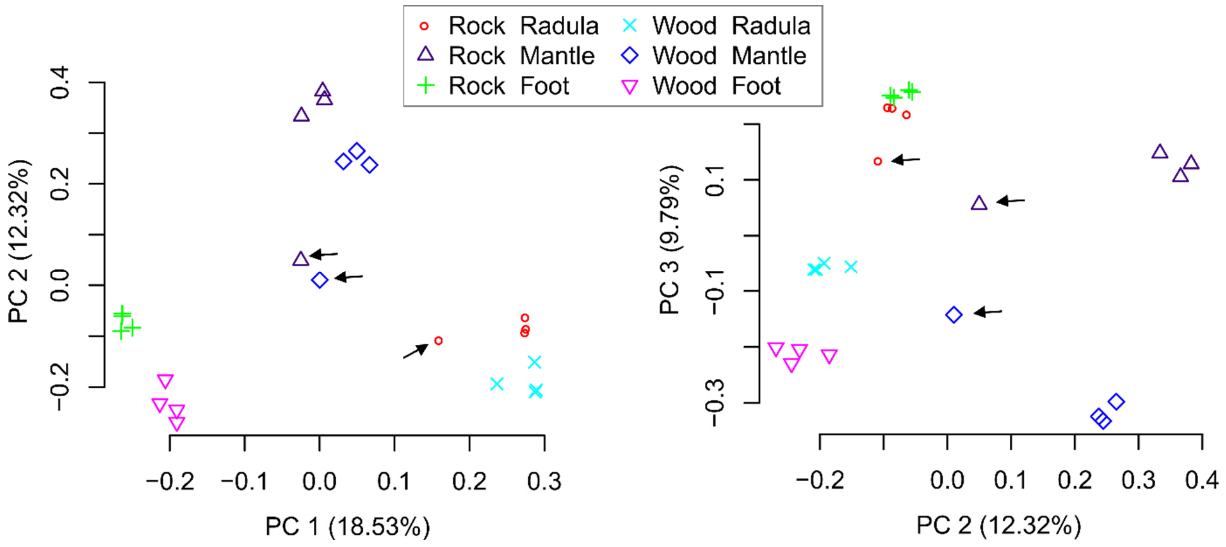


**Suppl. figure 3:** Principal component analyses of gene expression before filtering including outlier samples. Tissue samples that led us to the decision to exclude samples of pool1 of both ecomorphs are marked with black arrows.

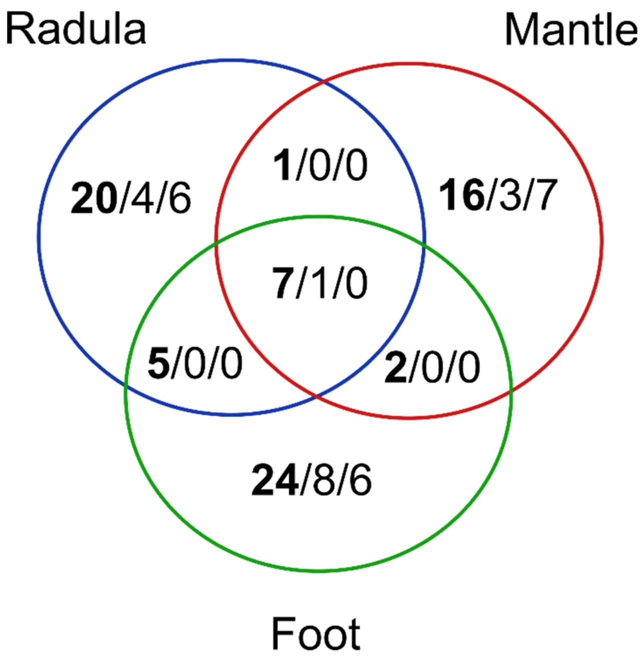


**Suppl. figure 4:** Venn graph illustrating the presence of alternatively fixed SNPs in transcripts of genes that are also differentially expressed ($p \leq 10^{-5}$) between at least one pair of identical tissues of both ecomorphs. The total number of SNPs in highly DE genes is shown first and in bold, followed by the number of synonymous and non-synonymous SNPs in these genes.

18

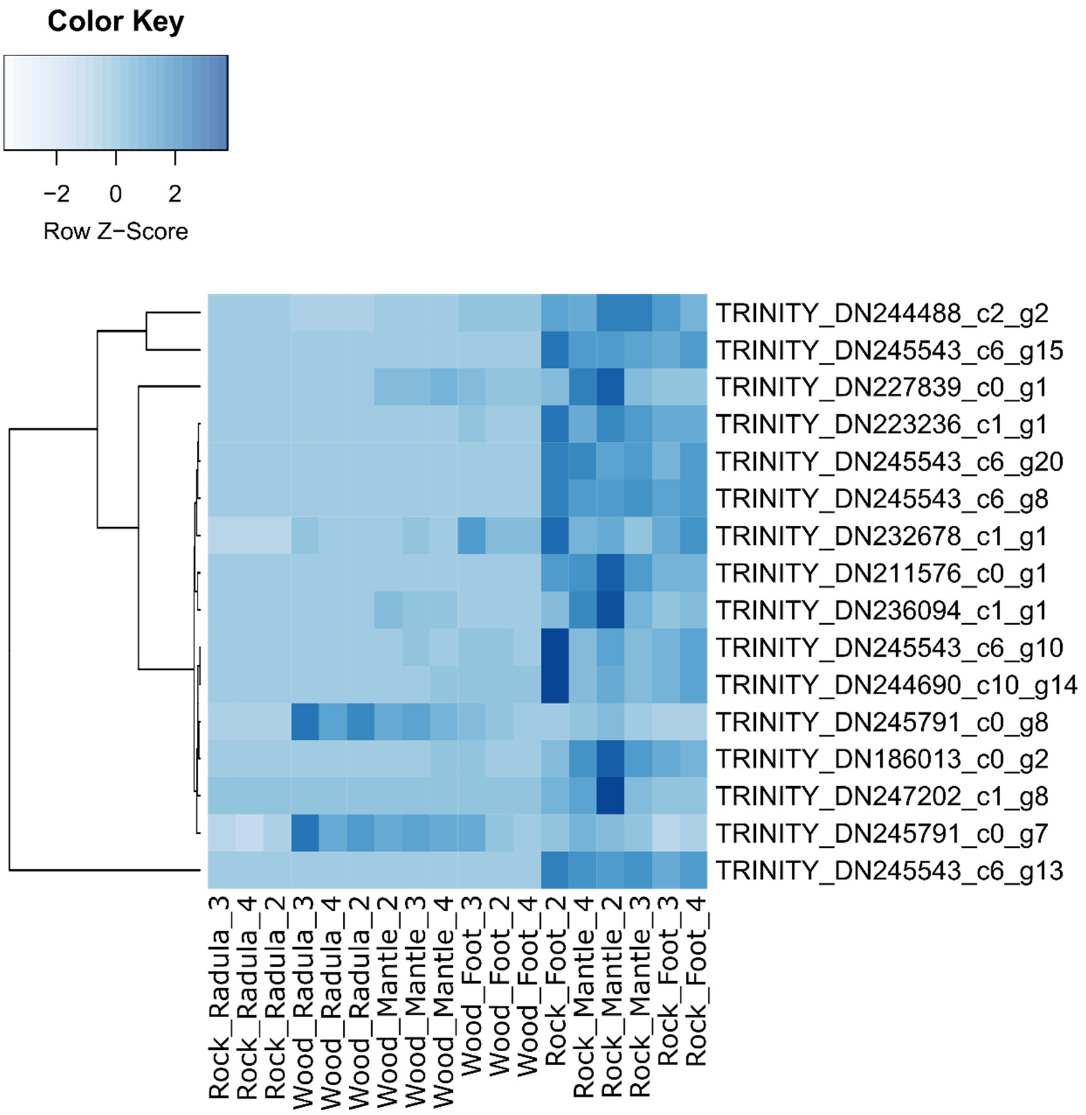


**Suppl. figure 5:** Gene expression of differentially expressed genes with the MF tetrapyrrole binding (GO:0046906). Hierarchical clustering heatmap of tetrapyrrole binding genes that were enriched among differentially expressed genes between ecomorphs. Samples and genes with similar expression cluster together. Heatmap is colored according to the row-wise z-score, which means that genes overexpressed in a certain sample relative to the other samples are colored dark blue, while underexpressed genes are colored light blue in the heatmap.

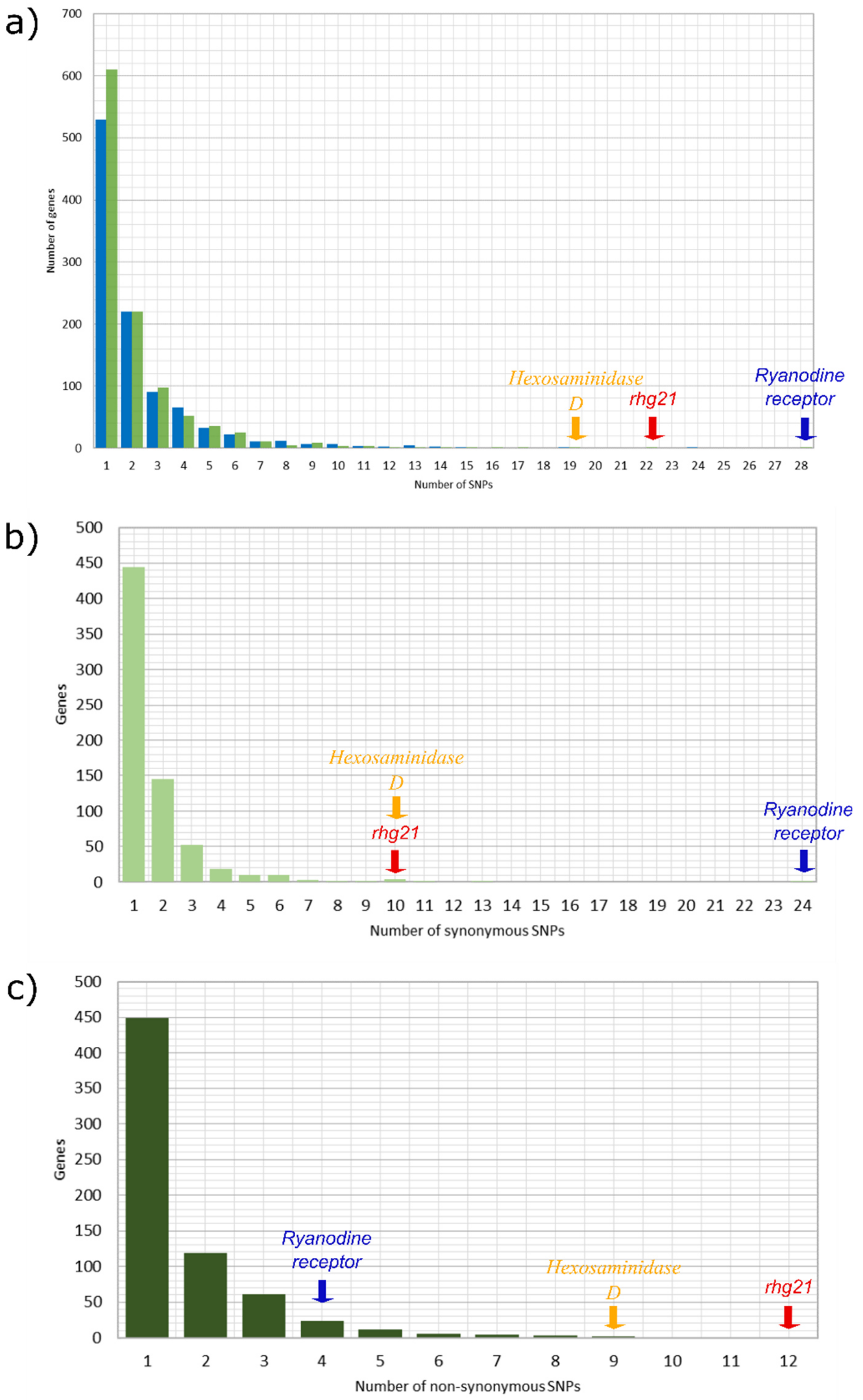


**Suppl. figure 6:** Distributions of alternatively fixed SNP numbers per transcript. a) Numbers of alternatively fixed SNPs per transcript inside (green) and outside (blue) of ORFs and the number of b) synonymous and c) non-synonymous alternatively fixed SNPs per gene are shown. Colored arrows indicate the three genes with the highest number of SNPs inside their ORFs in a) and show the number of synonymous and non-synonymous SNPs in b) and c), respectively. Only transcripts that carry at least one alternatively fixed SNP are included in this figure.

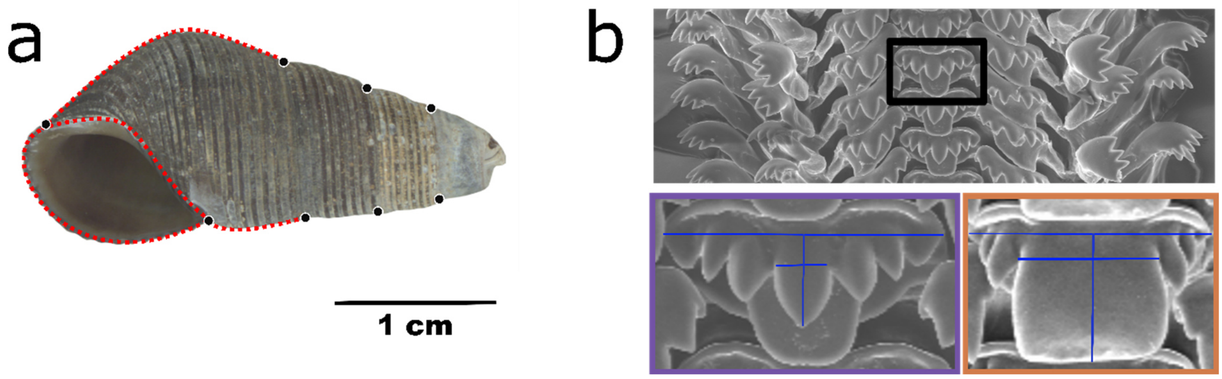


**Suppl. figure 7:** Morphological characterization of *T. sarasinorum* shell and radula. a) Landmarks (black) and semi-landmarks (red) were positioned on digital photographs of *T. sarasinorum* shells from individuals collected from wood and rock substrates at Loeha Island. All semilandmarks consisted of 10 sliding landmarks with the exception of the shortest stretch from the top of the aperture to the top of the first whorl (most bottom right semilandmark). b) Length measurements (blue lines) of the rachis (black box) and its central rachis denticle were used to characterize radula shape of *T. sarasinorum* wood (purple) and rock (orange) ecomorphs.

**Suppl. table 1** Samples and number of *Tylomelania sarasinorum* specimens from different substrates of which radulae were inspected for changes along the radula using SEM.

| Acc.-No. | N radulae studied | Substrate |
|---|---|---|
| ZMB Moll. 190370a | 12 | wood |
| ZMB Moll. 190370b | 9 | rocks |
| ZMB Moll. 190371a | 13 | wood |
| ZMB Moll. 190371b | 13 | rocks |
| ZMB Moll. 190372a | 2 | wood |
| ZMB Moll. 190372b | 2 | rocks |
| ZMB Moll. 190373a | 11 | wood |
| ZMB Moll. 190373b | 12 | rocks |
| ZMB Moll. 190374a | 1 | wood |
| ZMB Moll. 190375a | 25 | wood |
| ZMB Moll. 190375b | 10 | rocks |
| ZMB Moll. 190376a | 12 | rocks |
| ZMB Moll. 190376b | 12 | wood |
| ZMB Moll. 190377a | 19 | rocks |
| ZMB Moll. 190377b | 6 | wood |
| ZMB Moll. 190378a | 12 | wood |
| ZMB Moll. 190378b | 12 | rocks |
| ZMB Moll. 190379a | 12 | wood |
| ZMB Moll. 190379b | 6 | rocks |
| ZMB Moll. 190380a | 8 | wood |
| ZMB Moll. 190380b | 12 | rocks |
| ZMB Moll. 190381a | 6 | wood |
| ZMB Moll. 190381b | 12 | rocks |
| ZMB Moll. 190382a | 2 | wood |
| ZMB Moll. 190382b | 8 | rocks |
| ZMB Moll. 190383a | 10 | wood |
| ZMB Moll. 190383b | 12 | rocks |
| ZMB Moll. 190384a | 12 | wood |
| ZMB Moll. 190384b | 12 | rocks |
| ZMB Moll. 190385a | 5 | rocks |
| ZMB Moll. 190385b | 6 | wood |
| ZMB Moll. 190703b | 8 | wood |
| ZMB Moll. 190703c | 3 | leaves |
| ZMB Moll. 190724 | 5 | wood |
| ZMB Moll. 190729a | 6 | rocks |
| ZMB Moll. 190729b | 6 | wood |
| ZMB Moll. 190732a | 6 | rocks |
| ZMB Moll. 190732b | 6 | wood |
| ZMB Moll. 190740a | 2 | rocks |
| ZMB Moll. 190740b | 6 | wood |
| ZMB Moll. 190742a | 2 | rocks |

| | | |
|---|---|---|
| ZMB Moll. 190742b | 5 | wood |
| ZMB Moll. 190753a | 6 | rocks |
| ZMB Moll. 190753b | 5 | wood |
| ZMB Moll. 190763a | 6 | rocks |
| ZMB Moll. 190763b | 8 | wood |
| ZMB Moll. 191046a | 3 | rocks |
| ZMB Moll. 191046b | 3 | wood |
| ZMB Moll. 191051 | 1 | mud |
| ZMB Moll. 191052 | 1 | wood |
| ZMB Moll. 191059a | 3 | rocks |
| ZMB Moll. 191059b | 4 | wood |
| ZMB Moll. 191531a | 8 | rocks |
| ZMB Moll. 191531b | 6 | wood |
| ZMB Moll. 191532a | 6 | rocks |
| ZMB Moll. 191532b | 6 | wood |
| ZMB Moll. 191533a | 6 | rocks |
| ZMB Moll. 191533b | 6 | wood |
| ZMB Moll. 191534 | 12 | rocks |
| ZMB Moll. 191535 | 5 | rocks |
| ZMB Moll. 191536a | 13 | rocks |
| ZMB Moll. 191536b | 6 | wood |
| ZMB Moll. 191537a | 3 | rocks |
| ZMB Moll. 191537b | 5 | wood |
| ZMB Moll. 191538a | 3 | rocks |
| ZMB Moll. 191538b | 6 | wood |
| ZMB Moll. 191539a | 2 | rocks |
| ZMB Moll. 191539b | 2 | wood |

**Suppl. table 2** Principle components of shell shape and the proportion of variance explained.

| | Standard deviation | Proportion of variance | Cumulative variance |
|---|---|---|---|
| **PC1** | 0.02157 | 0.42405 | 0.42405 |
| **PC2** | 0.01138 | 0.11802 | 0.54207 |
| **PC3** | 0.01043 | 0.09905 | 0.64112 |
| **PC4** | 0.008205 | 0.061340 | 0.702460 |
| **PC5** | 0.00757 | 0.05221 | 0.75467 |
| **PC6** | 0.006746 | 0.041470 | 0.796140 |
| **PC7** | 0.006332 | 0.036540 | 0.832680 |
| **PC8** | 0.005894 | 0.031650 | 0.864330 |
| **PC9** | 0.004971 | 0.022520 | 0.886850 |
| **PC10** | 0.004803 | 0.021020 | 0.907870 |
| **PC11** | 0.003991 | 0.014510 | 0.922380 |
| **PC12** | 0.003837 | 0.013420 | 0.935800 |
| **PC13** | 0.003287 | 0.009850 | 0.945650 |
| **PC14** | 0.003046 | 0.008450 | 0.954100 |
| **PC15** | 0.002852 | 0.007410 | 0.961520 |

**Suppl. table 3** T-test results for shell size and PCs of shape as well as PC1 of radula shape. Significant differences between ecomorphs are indicated by bold p-values.

| Trait | | t | df | *p* |
|---|---|---|---|---|
| Shell | PC1 | 0.95 | 33.47 | 0.351 |
| Shell | PC2 | 5.67 | 33.01 | **0.000** |
| Shell | PC3 | 0.54 | 28.48 | 0.590 |
| Shell | PC4 | 0.29 | 34.99 | 0.774 |
| Shell | PC5 | -0.004 | 32.75 | 0.997 |
| Shell | PC6 | -1.11 | 32.19 | 0.275 |
| Shell | Size | -1.30 | 33.31 | 0.203 |
| Radulae | PC1 | -5.22 | 93.18 | **0.000** |

**Suppl. table 4** Enriched gene ontologies in genes with alternatively fixed non-synonymous SNPs and differentially expressed genes between identical tissues of both ecomorphs.

| Gene set | GO | Description | Log10 (p-value) |
|---|---|---|---|
| DE genes between ecomorphs | **BP** | | |
| | - | - | - |
| | **CC** | | |
| | - | - | - |
| | **MF** | | |
| | GO:0004497 | monooxygenase activity | -4.97 |
| | GO:0016712 | oxidoreductase activity acting on paired donors with incorporation or reduction of molecular oxygen reduced flavin or flavoprotein as one donor and incorporation of one atom of oxygen | -4.83 |
| | GO:0016705 | oxidoreductase activity acting on paired donors with incorporation or reduction of molecular oxygen | -4.18 |
| | GO:0005506 | iron ion binding | -4.18 |
| | GO:0046906 | tetrapyrrole binding | -3.64 |
| | GO:0016491 | oxidoreductase activity | -2.70 |
| | GO:0030246 | carbohydrate binding | -2.70 |
| | GO:0020037 | heme binding | -2.48 |
| Genes with alternatively fixed non-synonymous SNPs | **BP** | | |
| | GO:0008150 | Biological process | -2.35 |
| | **CC** | | |
| | GO:0005575 | Cellular component | -2.77 |
| | **MF** | | |
| | - | - | - |

**Suppl. table 5** Number of paired-end reads before and after quality filtering.

| Sample | # mio raw pe reads | # mio quality filtered pe reads |
|---|---|---|
| Pool1 Wood Mantle | 52.63 | 45.47 |
| Pool1 Wood Radula | 136.52 | 121.5 |
| Pool1 Wood Foot | 44.13 | 38.51 |
| Pool2 Wood Mantle | 37.11 | 32.82 |
| Pool2 Wood Radula | 29.14 | 26.23 |
| Pool2 Wood Foot | 41.69 | 36.1 |
| Pool3 Wood Mantle | 32.5 | 28.31 |
| Pool3 Wood Radula | 41.36 | 36.96 |
| Pool3 Wood Foot | 47.33 | 41.52 |
| Pool4 Wood Mantle | 34.33 | 29.69 |
| Pool4 Wood Radula | 27.21 | 24.26 |
| Pool4 Wood Foot | 37.29 | 31.92 |
| Pool1 Rock Mantle | 54.19 | 47.07 |
| Pool1 Rock Radula | 65.64 | 58.16 |
| Pool1 Rock Foot | 45.07 | 39.15 |
| Pool2 Rock Mantle | 37.07 | 33.1 |
| Pool2 Rock Radula | 38.73 | 34.45 |
| Pool2 Rock Foot | 39.21 | 34.3 |
| Pool3 Rock Mantle | 30.62 | 27.24 |
| Pool3 Rock Radula | 39.71 | 35.49 |
| Pool3 Rock Foot | 38.67 | 33.72 |
| Pool4 Rock Mantle | 37.6 | 33.29 |
| Pool4 Rock Radula | 38.98 | 34.8 |
| Pool4 Rock Foot | 42.46 | 37.09 |
| **Sum** | 1069.19 | 941.15 |
